# Biological validation of fecal corticosterone metabolites as a non-invasive stress assessment in translocated California valley quail (*Callipepla californica*)

**DOI:** 10.1101/2023.10.26.564168

**Authors:** Sarah A. Currier, Jeffrey G. Whitt, Kelly S. Reyna

## Abstract

U.S. quail species are vulnerable to population declines as a result climate change, habitat loss, and habitat fragmentation; all of which can induce physiological stress. Additionally, population restoration techniques (PRTs), like translocations, also induce stress. Traditional stress assessments include capturing and handling birds to extract blood, methods which are inherently stressful and can compound stress analyses. However, the stress hormone corticosterone is metabolized from the blood and excreted in feces as fecal corticosterone metabolites (FCMs). FCMs have been used as a non-invasive measurement of stress in a variety of species, but must be validated for each species. The objective of this study was to biologically validate the use of FCMs as a non-invasive measurement of stress-hormone levels in California valley quail (*Callipepla californica*). Reference and treatment quail were acclimated for 3 wks in an outdoor aviary. Subsequently, treatment quail were subjected to a simulated, 48-h translocation, a common and stressful PRT. Fecal samples were collected every 4 h and processed using an enzyme immunoassay. Mean FCM concentrations of treatment quail (41.50 ±16.13 ng/g) were higher than reference FCM concentrations (24.07 ±10.4 ng/g). These results biologically validate the use of FCMs as a non-invasive method to assess stress hormone levels in California valley quail, demonstrate diurnal variation in quail stress levels, and confirms that quail translocations are a stressful PRT. Ultimately, this research validates a new non-invasive tool for stress measurement to advance quail research, management, and conservation.

**Lay summary:** This study biologically validates the use of fecal corticosterone metabolites as a non-invasive method for detecting stress in quail, demonstrates diurnal variation in quail stress levels, confirms that translocations elevate stress which likely impacts success, and establishes a new non-invasive tool for stress measurement in quail research, management, and conservation.

## Introduction

Stress is a factor often overlooked in U.S. quail population declines and population restoration techniques. U.S. quail populations are vulnerable to decline as a result of climate change (Guthery *et al*., 2000; Tanner *et al*., 2017; Wilsey *et al*., 2019), habitat loss, and habitat fragmentation (Brennan and Covington, 1994; Hernández *et al*., 2013; Pope and Heekin, 2017); all of which are stressful to quail as indicated by increased levels of blood corticosterone, (CORT, 17-deoxycortisol), a steroid hormone produced in the cortex of the adrenal gland (Creel, 2001; Suorsa *et al*., 2003; Tempel and Gutiérrez, 2004; Cockrem, 2007, 2013; Sheriff *et al*., 2011; Wingfield *et al*., 2017; Palme, 2019; van Vliet *et al*., 2020). For example, elevated CORT levels have been observed in birds as a result of anthropogenic disturbances (Thiel *et al*., 2011; Jankowski *et al*., 2014; Arlettaz *et al*., 2015) and extreme climatic events (Wingfield *et al*., 2017), which are expected to increase in frequency and severity due to global climate change (Seneviratne *et al*., 2012; IPCC, 2021). Stress from climate change can have synergistic effects, multiplying the deleterious impacts of other stressors (Felton *et al*., 2009; Şekercioğlu *et al*., 2012).

Elevated stress levels can directly influence bird populations by altering foraging, predator avoidance, fledging, and other behaviors (Cyr and Romero, 2007; Crespi *et al*., 2013; Wingfield *et al*., 2017). Elevated CORT levels in birds can also lead to increased predator encounters (Stephens *et al*., 2004), which further increases stress levels (Clinchy *et al*., 2013). Extreme stress can induce a type 1 allostatic overload, where birds decrease or eliminate behaviors (e.g., reproduction) that are not immediately necessary for survival (McEwen and Wingfield, 2003) by increasing gonadotropin-inhibitory hormone, reducing gonadotropin-releasing hormone (GnRH), and displaying a reduced sensitivity to GnRH and luteinizing hormone (Sapolsky *et al*., 2000; Wingfield and Sapolsky, 2003; Dickens and Bentley, 2014); though the mechanisms are not completely understood (Wingfield and Ramenofsky, 2011; Lattin *et al*., 2016). For example, exposing zebra finches *(Taeniopygia guttata*) to stress stimuli increased CORT levels and decreased testosterone, resulting in reduced reproductive behavior (Lynn *et al*., 2010). Capturing and holding wild European starlings (*Sturnus vulgaris*) blocked a natural shift in the hypothalamic-pituitary-adrenal (HPA) axis from early breeding (gonadal development) to active breeding activity, which inhibited breeding (Dickens and Bentley, 2014). The resultant reduction in reproduction in birds contributes to population declines.

Population restoration techniques (PRTs) can also increase CORT levels. U.S. quail species are economically important gamebirds (Johnson *et al*., 2012; Wszola *et al*., 2020) and part of a $3.7 billion per year upland gamebird hunting industry (Sportsmen’s Alliance, 2021). For over 150 years, a variety of PRTs like translocations have been used in attempts to bolster populations (Hernández *et al*., 2013; Gomez and Reyna, 2017; Whitt *et al*., 2017). Quail Translocations in general are rarely successful (Perez *et al*., 2002; Scott *et al*., 2013; Whitt *et al*., 2017), especially when used for reintroduction (Martin *et al*., 2017). Although these PRTs have rarely resulted in long-term success, there is increasing interest in translocation of wild quail (Stephenson *et al*., 2011; Martin *et al*., 2017; Sisson *et al*., 2017). Numerous hypotheses for the low success rate have been proposed (Sokos *et al*., 2008; Martin *et al*., 2017), but little is known about how stress influences translocation success.

Translocation consists of capturing, handling, holding, transporting, and releasing quail to a novel site, each of which can result in acute and chronic stress, independent of other processes (Buchanan, 2000; Fazio and Ferlazzo, 2003; Jones *et al*., 2005; Dickens *et al*., 2009, 2010; Wingfield and Romero, 2011; Batson *et al*., 2017; Martin *et al*., 2017). For example, short translocations may induce acute stress; however, the duration of a quail translocation is typically 1-3 days, which increases the occurrence of chronic stress (Dickens *et al*., 2010).

Chronic stress can disrupt the fight or flight response as a result in downregulation of hormonal responses to acute and chronic stressors, (Romero and Butler, 2007; Dickens *et al*., 2009; Martin *et al*., 2017). For example, European starlings caught in the wild and placed into captivity lost the ability to produce a fight or flight response when presented with a stress stimulus (Dickens and Romero, 2009). In addition, when the fight or flight response was chronically stimulated it negatively impacted cardiovascular health, leading to hypertension due to surplus catecholamine exposure (Rupp, 1999). Both chronic and acute stress responses can inhibit translocated quail survival (Armstrong and Seddon, 2008; Dickens *et al*., 2010; Breed *et al*., 2019), and lead to failure of the translocation by increasing the susceptibility to predation (Dickens *et al*., 2010; Martin *et al*., 2017).

Weight loss has been observed when translocating wild birds to novel environments (Rich and Romero, 2005; Dickens *et al*., 2009; Fischer *et al*., 2018), and is a symptom of chronic stress. For example, European starlings lost 5–15% of their body weight after being exposed to a 14-d regime of multiple stressors during a translocation (Awerman and Romero, 2010).

California valley quail (*C. californica*) exhibited a mean weight loss of 14.3% when translocated from Idaho to Texas (Reyna *et al*., 2020). This is important because body mass can influence survival, and the overall success of translocations. For example, Cirl buntings (*Emberiza cirlus*) with a higher body weight at capture were more likely to survive in their new habitat than those with lower body weights (Fountain *et al*., 2017). Reducing the amount of stress during a translocation could reduce weight loss and increase post-release survival (Warwick *et al*., 2006).

The traditional method of measuring stress in birds requires capturing, handling, and extracting blood samples; inherently stressful events (Arnold *et al*., 2008), followed with measuring blood CORT levels. It is assumed that if this process occurs within 3 min of initial handling, CORT levels will be indicative of the physiological condition of the bird prior and will not represent the elevated stress incurred during the procedure (Littin and Cockrem, 2001; Cockrem and Silverin, 2002). However, CORT levels quickly rise during handling and peak within 15–30 min. In addition, handling associated with blood extractions adds to the cumulative stress experienced by the bird, and nearby birds, independent of the extraction.

Steroid hormones, like CORT, can also be measured from urine, feces, hair, and feather samples (Bortolotti *et al*., 2008; Sheriff *et al*., 2011). One effective, non-invasive, and increasingly popular method to evaluate stress is by measuring fecal corticosterone metabolites (FCMs) (Romero and Remage-Healey, 2000; Tempel and Gutiérrez, 2004; Möstl *et al*., 2005; Dickens *et al*., 2009; Sheriff *et al*., 2010). Measuring FCMs as an assessment of stress has been successful in a wide range of avian studies (Washburn *et al*., 2003; Fletcher *et al*., 2018; Sokół and Koziatek-Sadłowska, 2020). Excretion of FCMs varies between species, as does the suitability of different immunoassays to accurately detect CORT, therefore methods must be validated for each species (Palme *et al*., 1996; Wasser *et al*., 2000; Palme, 2019). Physiological validation of measuring FCM concentrations requires injecting a known quantity of CORT into an animal (Touma and Palme, 2005), an invasive and stressful procedure (Palme, 2019; Mohlman *et al*., 2020). Biological validation of measuring FCM concentrations is a non-invasive procedure where feces is collected before and after an event that increases HPA activity for an extended time, resulting in increased CORT (Touma and Palme, 2005; Palme, 2019).

The goal of this study was to biologically validate the use of FCMs as a non-invasive measurement of stress in California valley quail using a simulated translocation as the stressor. This is the first biological validation of FCMs as a tool to evaluate stress in a new world quail species. Mohlman et al. (2020) performed a physiological validation of FCMs with northern bobwhite (*Colinus virginianus*), a close relative, but were unsuccessful at biological validation.

## Materials and methods

Flight-ready captive-reared valley quail (*N* = 5 females, *N* = 8 males), 16–24 wks in age, were acquired from a breeder (ZKD Game Birds, West Point, Texas, USA) in October, 2020. Quail were housed and acclimated in an outdoor aviary (Quail Hotel, Fannin Fabrication, Bonham, TX, USA) for 3 wks. Birds were provided with gamebird feed (Gamebird Starter and Grower, M-G Inc., Weimar, TX, USA) and water *ad libitum,* and enrichments including social and environmental stimuli (Hawkins et al. 2001). This mimicked the time and conditions in which quail were held prior to the first shipment of trapped birds during a 2019 translocation (Reyna *et al*., 2020). Acclimation was assumed when birds exhibited typical behavior (e.g., eating, drinking, roosting, grooming; Reyna and Newman, 2018).

To biologically validate FCMs in valley quail, reference (low stress) fecal samples were obtained from acclimated valley quail. The floor of the aviary was lined with a plastic sheet and fecal samples were collected daily. Once collected, individual fecal samples were placed into a plastic storage bag (Ziploc freezer quart, S. C. Johnson & Son, Inc., Racine, WI) and labeled with the date, time, and sex of the bird. Temperature, humidity, and presence of direct sunlight were recorded on the plastic storage bag, since these factors can degrade hormone detection probability (Shipley *et al*., 2019). Immediately after collection and labeling, samples were stored in a freezer at -20 ℃ until FCM analysis (Wasser *et al*., 1988; Messmann *et al*., 1999; Khan *et al*., 2002).

For biological validation, quail were transported to the animal care facility at Texas A&M University-Commerce to undergo a simulated 48-h translocation, a stressful event (Martin et al. 2017). Each quail was given a unique leg band, and had their age, sex, and weight recorded. Quail were held individually in research-approved breeding pens located in a temperature and light-controlled room with a 12L:12D photoperiod. To ensure accuracy and reduce contamination, fecal samples were collected from the underlying waste pan trays 4–5 times daily (Millspaugh and Washburn, 2004; Mohlman *et al*., 2020). The frequency of feces collection ensured a representative sample, since FCMs may fluctuate diurnally (Breuner *et al*., 1999).

Individual fecal samples were placed in a plastic storage bag and labeled with the bird ID number, sex, time, and date of collection. Steroid hormone concentrations can be affected by storage duration (Millspaugh and Washburn, 2004) and bacteria (Lexen *et al*., 2008). Therefore, high stress fecal samples were immediately stored in a freezer (-20 ℃) until FCM analysis (Wasser *et al*., 1988; Messmann *et al*., 1999).

FCM concentrations of reference and stressed quail fecal samples were measured using an Enzyme Immunoassay (EIA) Kit (Arbor Assays, Ann Arbor, MI, USA). The kit was validated for use on CORT extracted from dry fecal extracts on a multi-species design (Jackson, 2015; Fletcher *et al*., 2018), and included detailed instructions for extracting CORT from fecal samples and analyzing FCM concentrations. To determine the assays suitability for valley quail, 10 fecal extracts were pooled from the valley quail simulated translocation and a serial dilution (*N* = 6 dilutions) was performed. FCM concentrations from the pooled fecal extracts were parallel to the slope of the standard curve, verifying suitability for valley quail. The sensitivity of the EIA was determined to be 18.19 pg/ml. Optical density was calculated using a Synergy LX Multi-mode microplate reader (Bio Tek Instruments, USA), and entered into MyAssay web (https://www.myassay.com), to calculate FCM concentration in pg/ml. FCM concentrations reported in pg/ml were converted to ng/g for comparison to other studies.

All animals were handled in accordance with procedures outlined in the Guide for the Care and Use of Laboratory Animals (National Research Council, 2010), and Texas A&M University-Commerce Animal Use Protocol P20-013.

### Statistical Analysis

All statistical analyses were conducted in R (version 4.0.2, R Foundation, Austria). Shapiro-Wilk’s statistical test was used to check for normality (Zar, 1996). Welch’s 2-sample t-test was used to calculate the p-value, mean, and 95% confidence interval between 2 variables with normal distributions. Mann-Whitney *U*-test was used to compare 2 variables with non-normal distributions. Kruskal-Wallis test was used to compare multiple variables with non-normal distributions, followed by Dunn’s test with Bonferroni correction for pairwise comparisons.

Correlations were tested using Pearson’s product-moment correlation with Bonferroni correction (Zar, 1996). All weight data are presented as mean ± SD. Results were significant at alpha <0.05.

## RESULTS

Mean ±SD FCM concentrations of valley quail during the simulated translocation were 36.21 ±18.56 ng/g, and higher than reference FCM concentrations (24.07 ±10.4 ng/g; *N* = 13 quail; Mann-Whitney *U*, *Z* = 4.20, *P* < 0.001; Fig. 1). Mean 48-hour weight loss was 17.8 ±6.3 g, or 12 ±4.4% of initial body mass.

**Fig 1.**
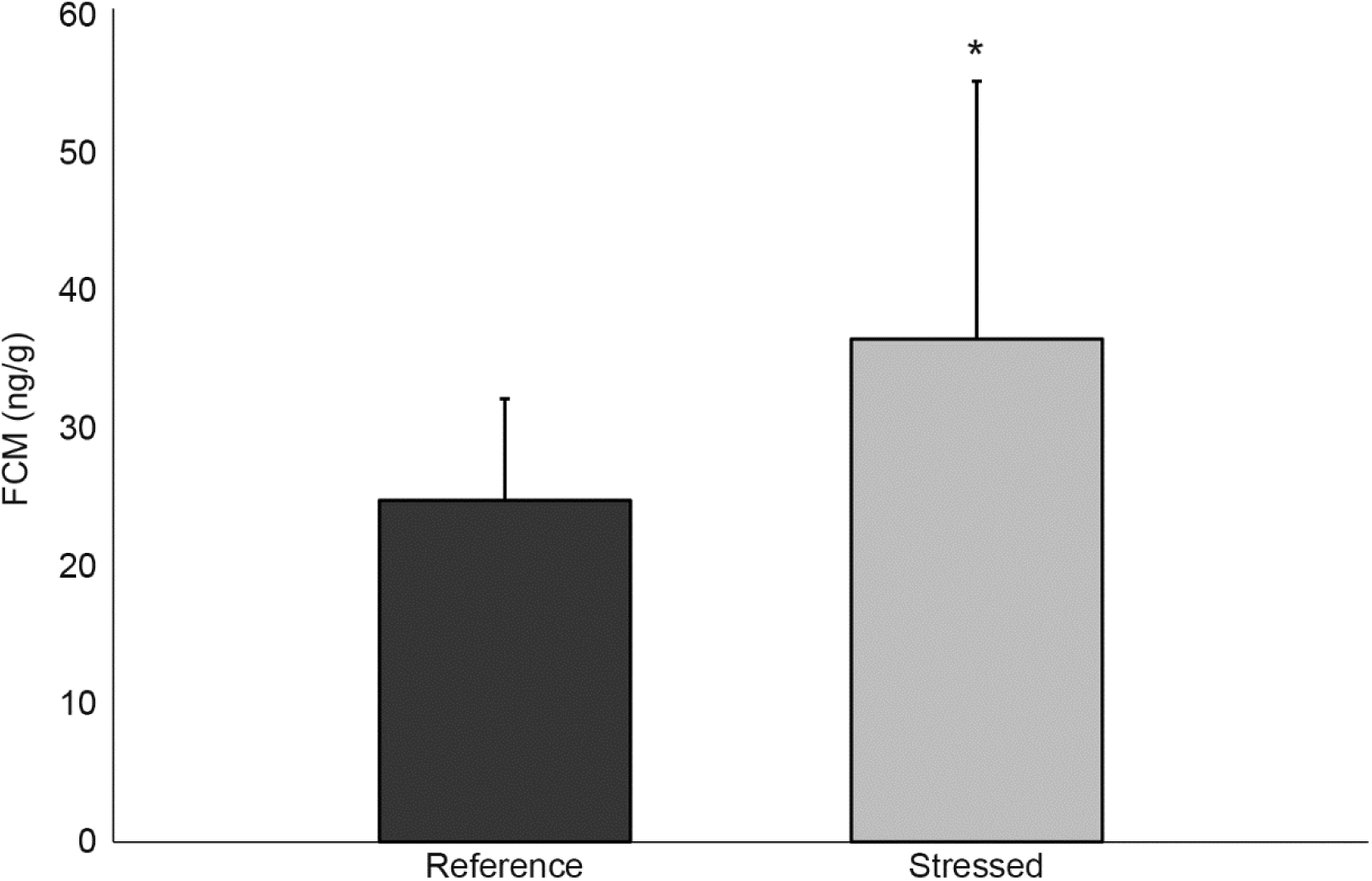
Biological validation of using fecal corticosterone metabolites in translocated California valley quail. Fecal corticosterone metabolites (FCM) concentrations (mean ±SD) were extracted from scat collected from acclimated quail (N=13) prior to (Reference) and during a 48-hr simulated translocation (Stressed). FCM concentrations from stressed birds were higher than reference FCM concentrations (Mann-Whitney *U*, *Z* = 4.20, *P* < 0.001). Asterisk indicates statistical difference.

Valley quail with higher initial body mass prior to the simulated translocation experienced lower mean FCM concentrations (r^2^ = 0.14, *P* < 0.001), and lower percentage of initial mass lost (r^2^ = 0.13, *P* < 0.001) during the simulated translocation. FCM concentrations positively correlated with percent of initial body mass loss (r^2^ = 0.27, *P* < 0.001). No difference in FCM concentrations (Mann-Whitney *U*, *Z* = 0.15, *P =* 0.44), total mass lost (Welch’s t-test, *P* = 0.58), or percentage of initial mass lost (Welch’s t-test, *P* = 0.29) was observed based on sex.

Valley quail FCMs varied diurnally during the simulated translocation (Fig. 2), with higher concentrations from 10:00–18:00 and lower concentrations from 20:00–06:00 (Kruskal-Wallis, H = 62.3, *P* < 0.001). Mean FCM concentrations were not different between the first 24 h and last 24 h (Mann-Whitney *U*, *Z* = 1.86, *P* = 0.32).

**Fig 2.**
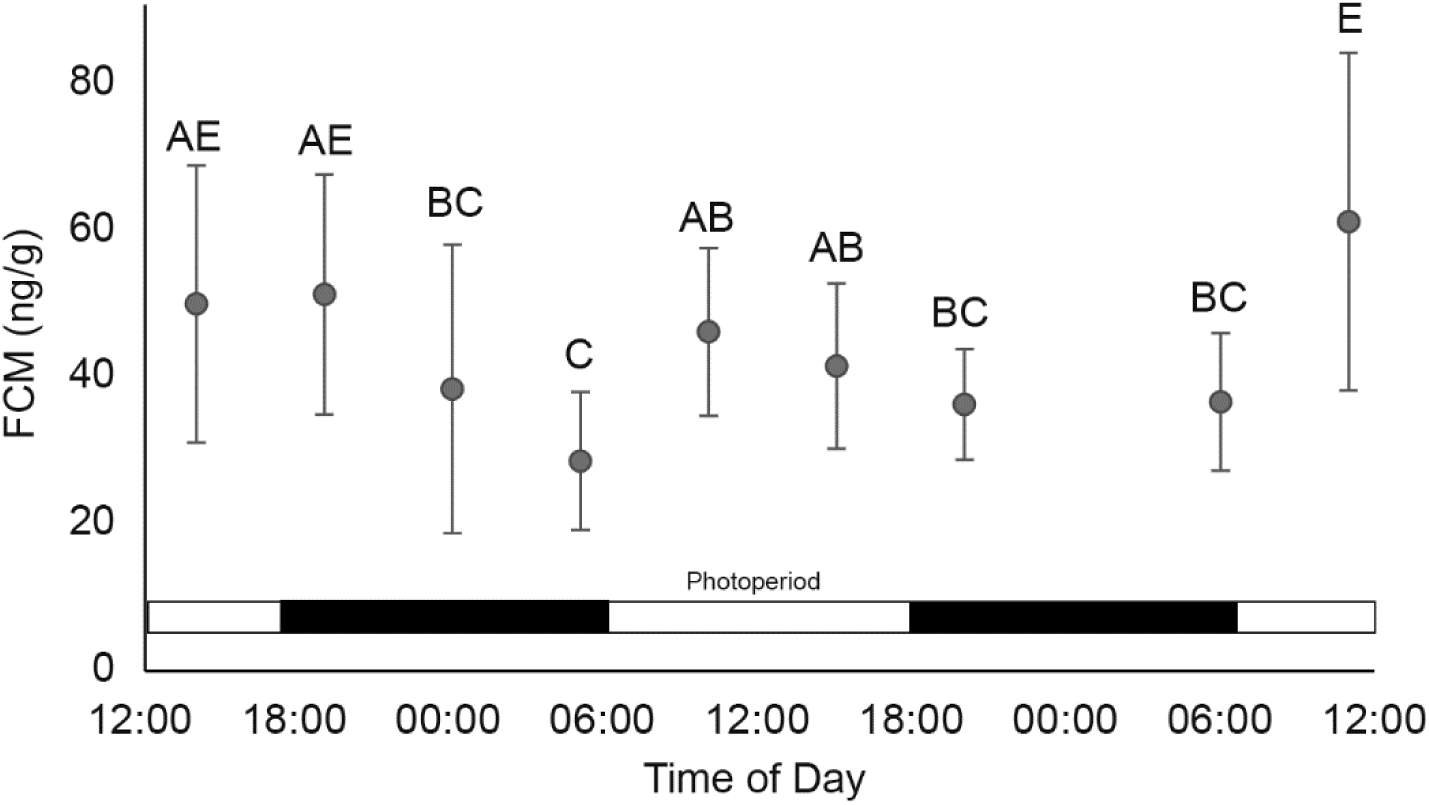
Diurnal variation in fecal corticosterone metabolite (FCM) concentrations from California valley quail during a simulated 48-h translocation. FCM concentrations correlated with photoperiod (12L:12D) and quail circadian rhythm. Letters indicate statistical groupings.

## Discussion

This study was the first biological validation of fecal corticosterone metabolites as a non-invasive assessment of stress for a new world quail species. When exposed to a simulated translocation, valley quail experienced a 72.4 % increase in FCMs compared to reference concentrations, indicating that this protocol successfully detected an increase in FCMs during a stressful event. These results are comparable to the 73% increase in FCMs recorded in northern bobwhites during a physiological validation (Mohlman *et al*., 2020). However, the increase in FCM levels is lower than that recorded in many other avian species. This isn’t surprising because excretions of metabolites can vary between species (Palme *et al*., 1996; Wasser *et al*., 2000; Palme, 2019). For example. experimentally stressed European starlings showed a ∼100% increase in FCMs (Cyr and Romero, 2008). Following capture and holding for a veterinary examination, African penguins (*Spheniscus demersus*) showed a 155–349% increase in FCMs.

Wild Dickcissels (*Spiza americana*) experienced a 1,700% increase in FCM concentrations within 24 h of having leg harness transmitters attached (Suedkamp Wells *et al*., 2003). Valley quail and northern bobwhite quail are both new world quail and their similar stress response during validation, and differences from other taxa were expected.

The correlation of weight loss with FCM levels further supports the association of chronic stress with weight loss in birds (Cyr and Romero, 2007; Dickens *et al*., 2009; Awerman and Romero, 2010). Quail in our study lost ∼12% of body weight which is consistent with weight loss in translocated starlings (5-15%) and wild valley quail (14.3 %) in an actual translocation of similar duration [60], indicating that translocation weight loss may primarily be a result of stress. Similar to our study, Reyna et al. (2020) reported that larger birds lost less weight during translocation. Larger birds may be healthier overall and future studies may benefit from considering a minimum weight for translocation.

By collecting fecal samples 4–5 times daily during the simulated translocation, we were able to capture diurnal variation in FCM concentrations. While not previously observed in new world quail (Mohlman *et al*., 2020), diurnal variation in corticosterone metabolites has been recorded in a variety of species (Florant and Weitzman, 1980; de Jong *et al*., 2001; Touma *et al*., 2004; Bosson *et al*., 2009; Sheriff *et al*., 2009), generally corresponding to the circadian rhythm of the animal. The diurnal variation in valley quail FCM concentrations was similar to variations observed in broiler chickens (de Jong *et al*., 2001), with lower concentrations at night and elevated concentrations during the day. This trend was expected since valley quail are active in the day and inactive during the night (Leopold, 1977).

This study verifies that translocations are stressful events for quail that result in significant weight loss; results that suggest stress influences translocation success. Chronic stress is associated with increased mortality (Cyr and Romero, 2007; Dickens *et al*., 2010), and animal weight at release is a predictor of survival (Fountain *et al*., 2017). Further, stress is a major driver of reproduction, which is one major indicator of translocation success (Griffith *et al*., 1989; Dickens *et al*., 2010). Future studies could benefit by focusing efforts on stress mitigation to increase long-term translocation success.

The use of FCM as a non-invasive measurement in new world quail could be extended beyond translocations to benefit quail conservation. There is considerable environmental and economic interest in conserving and restoring populations of quail and other gamebird species. Increased plasma CORT can be indicative of environmental disturbances and habitat-related metabolic challenges (Homyack, 2010; Shipley *et al*., 2022). For example, FCMs have been used as an indicator of declining habitat quality in greater sage-grouse (*Centrocercus urophasianus*; Rabon *et al*., 2021). FCMs have also been used to detect elevated CORT levels associated with extreme weather events, noise related to fossil fuel extraction (Cinto Mejia *et al*., 2019), and other anthropogenic disturbances (Thiel *et al*., 2008, 2011; Blickley *et al*., 2012; Arlettaz *et al*., 2015; Formenti *et al*., 2015). Through the use of innovative field collection methods (Shipley *et al*., 2019), FCMs may be used to assess chronic stress levels in wild quail populations, evaluate responses to multiple stressors (e.g., extreme drought), and map stress levels across a landscape to identify focus areas for conservation (Rabon *et al*., 2021). Ultimately, the use of FCMs could alert quail biologists to the presence of environmental stressors before population demographics are impacted (Ellis *et al*., 2012). Clearly, FCMs have enormous potential as a non-invasive indicator of stress in quail and as a tool for improving quail research, restoration, management, and conservation.

## Acknowledgements

We thank C. Vandenberg for assisting with bird logistics, J. Delgado-Acevedo for project guidance, and Arbor Assays for technical support.

## Author Contributions

All authors contributed equally to this project and writing of this manuscript.

## Data Availability

The data underlying this article will be shared on reasonable requests to the corresponding author.

## References

Arlettaz R, Nusslé S, Baltic M, Vogel P, Palme R, Jenni-Eiermann S, Patthey P, Genoud M (2015) Disturbance of wildlife by outdoor winter recreation: allostatic stress response and altered activity–energy budgets. Ecol Appl 25: 1197–1212.

Armstrong DP, Seddon PJ (2008) Directions in reintroduction biology. Trends Ecol Evol 23: 20– 25.

Arnold JM, Oswald SA, Voigt CC, Palme R, Braasch A, Bauch C, Becker PH (2008) Taking the stress out of blood collection: comparison of field blood-sampling techniques for analysis of baseline corticosterone. J Avian Biol 39: 588–592.

Awerman JL, Romero LM (2010) Chronic psychological stress alters body weight and blood chemistry in European starlings (*Sturnus vulgaris*). Comp Biochem Physiol A Mol Integr Physiol 156: 136–142.

Batson WG, Gordon IJ, Fletcher DB, Portas TJ, Manning AD (2017) The effect of pre-release captivity on the stress physiology of a reintroduced population of wild eastern bettongs. J Zool 303: 311–319.

Blickley JL, Word KR, Krakauer AH, Phillips JL, Sells SN, Taff CC, Wingfield JC, Patricelli GL (2012) Experimental Chronic Noise Is Related to Elevated Fecal Corticosteroid Metabolites in Lekking Male Greater Sage-Grouse (*Centrocercus urophasianus*). PLoS One 7: e50462.

Bortolotti GR, Marchant TA, Blas J, German T (2008) Corticosterone in feathers is a long-term, integrated measure of avian stress physiology. Funct Ecol 22: 494–500.

Bosson CO, Palme R, Boonstra R (2009) Assessment of the Stress Response in Columbian Ground Squirrels: Laboratory and Field Validation of an Enzyme Immunoassay for Fecal Cortisol Metabolites. Physiol Biochem Zool 82: 291–301.

Breed D, Meyer LC, Steyl JC, Goddard A, Burroughs R, Kohn TA (2019) Conserving wildlife in a changing world: Understanding capture myopathy—A malignant outcome of stress during capture and translocation. Conserv Physiol 7: coz027.

Brennan LA, Covington WW (1994) Broad-Scale Population Declines in Four Species of North American Quail: An Examination of Possible Causes. In: Sustainable Ecological Systems: Implementing an Ecological Approach to Land Management. USDA Forest Service General Technical Report RM-247. pp 44–50.

Breuner CW, Wingfield JC, Romero LM (1999) Diel rhythms of basal and stress-induced corticosterone in a wild, seasonal vertebrate, Gambel’s white-crowned sparrow. J Exp Zool 284: 334–342.

Buchanan KL (2000) Stress and the evolution of condition-dependent signals. Trends Ecol Evol 15: 156–160.

Cinto Mejia E, McClure CJW, Barber JR (2019) Large-scale manipulation of the acoustic environment can alter the abundance of breeding birds: Evidence from a phantom natural gas field. J Appl Ecol 56: 2091–2101.

Clinchy M, Sheriff MJ, Zanette LY (2013) Predator-induced stress and the ecology of fear. Funct Ecol 27: 56–65.

Cockrem JF (2007) Stress, corticosterone responses and avian personalities. J Ornithol 148: 169–178.

Cockrem JF (2013) Corticosterone responses and personality in birds: Individual variation and the ability to cope with environmental changes due to climate change. Gen Comp Endocrinol 190: 156–163.

Cockrem JF, Silverin B (2002) Variation within and between Birds in Corticosterone Responses of Great Tits (Parus major). Gen Comp Endocrinol 125: 197–206.

Creel S (2001) Social dominance and stress hormones. Trends Ecol Evol 16: 491–497.

Crespi EJ, Williams TD, Jessop TS, Delehanty B (2013) Life history and the ecology of stress: how do glucocorticoid hormones influence life-history variation in animals? Funct Ecol 27: 93–106.

Cyr NE, Romero LM (2007) Chronic stress in free-living European starlings reduces corticosterone concentrations and reproductive success. Gen Comp Endocrinol 151: 82– 89.

Cyr NE, Romero LM (2008) Fecal glucocorticoid metabolites of experimentally stressed captive and free-living starlings: Implications for conservation research. Gen Comp Endocrinol 158: 20–28.

de Jong IC, van Voorst AS, Erkens JH, Ehlhardt DA, Blokhuis HJ (2001) Determination of the circadian rhythm in plasma corticosterone and catecholamine concentrations in growing broiler breeders using intravenous cannulation. Physiol Behav 74: 299–304.

Dickens MJ, Bentley GE (2014) Stress, captivity, and reproduction in a wild bird species. Horm Behav 66: 685–693.

Dickens MJ, Delehanty DJ, Romero LM (2009) Stress and translocation: alterations in the stress physiology of translocated birds. Proc R Soc Lond B Biol Sci 276: 2051–2056.

Dickens MJ, Delehanty DJ, Romero LM (2010) Stress: an inevitable component of animal translocation. Biol Conserv 143: 1329–1341.

Dickens MJ, Romero LM (2009) Wild European starlings *(Sturnus vulgaris*) adjust to captivity with sustained sympathetic nervous system drive and a reduced fight-or-flight response. Physiol Biochem Zool 82: 603–610.

Ellis RD, McWhorter TJ, Maron M (2012) Integrating landscape ecology and conservation physiology. Landscape Ecol 27: 1–12.

Fazio E, Ferlazzo A (2003) Evaluation of Stress During Transport. Vet Res Commun 27: 519– 524.

Felton A, Fischer J, Lindenmayer DB, Montague-Drake R, Lowe AR, Saunders D, Felton AM, Steffen W, Munro NT, Youngentob K, et al. (2009) Climate change, conservation and management: an assessment of the peer-reviewed scientific journal literature. Biodivers Conserv 18: 2243–2253.

Fischer CP, Wright-Lichter J, Romero LM (2018) Chronic stress and the introduction to captivity: how wild house sparrows (*Passer domesticus*) adjust to laboratory conditions. Gen Comp Endocrinol 259: 85–92.

Fletcher K, Xiong Y, Fletcher E, Gustafsson L (2018) Glucocorticoid response to both predictable and unpredictable challenges detected as corticosterone metabolites in collared flycatcher droppings. PloS One 13: e0209289.

Florant GL, Weitzman ED (1980) Diurnal and episodic pattern of plasma cortisol during fall and spring in young and old woodchucks (*Marmota monax*). Comp Biochem Physiol A Mol Integr Physiol 66: 575–581.

Formenti N, Viganó R, Bionda R, Ferrari N, Trogu T, Lanfranchi P, Palme R (2015) Increased hormonal stress reactions induced in an Alpine Black Grouse (*Tetrao tetrix*) population by winter sports. J Ornithol 156: 317–321.

Fountain K, Jeffs C, Croft S, Gregson J, Lister J, Evans A, Carter I, Chang YM, Sainsbury AW (2017) The influence of risk factors associated with captive rearing on post-release survival in translocated cirl buntings *Emberiza cirlus* in the UK. Oryx 51: 332–338.

Gomez LJ, Reyna KS (2017) An evaluation of northern bobwhite conservation research: a call for large-scale studies. Proceedings of the National Quail Symposium 8: 119–131.

Griffith B, Scott JM, Carpenter JW, Reed C (1989) Translocation as a species conservation tool: status and strategy. Science 245: 477–480.

Guthery FS, Forrester ND, Nolte KR, Cohen WE, Jr WPK (2000) Potential effects of global warming on quail populations. Proceedings of the National Quail Symposium 4: 198– 204.

Hernández F, Brennan LA, DeMaso SJ, Sands JP, Wester DB (2013) On reversing the northern bobwhite population decline: 20 years later. Wild Soc Bull 37: 177–188.

Homyack JA (2010) Evaluating habitat quality of vertebrates using conservation physiology tools. Wildl Res 37: 332–342.

Masson-Delmotte ZP, Pirani A, Connors SL, Péan C, Berger S, Caud N, Chen Y, Goldfarb L, Gomis MI et al. (2021) Climate change 2021: the physical science basis. In Masson-Delmotte, V., P. Zhai, A. Pirani, S.L. Connors, C. Pan, S. Berger, N. Caud, Y. Chen, L. Goldfarb, M.I. Gomis, M. Huang, K. Leitzell, E. Lonnoy, J.B.R. Matthews, T.K. Maycock, T. Waterfield, O. Yeleki, R. Yu, and B. Zhou (eds.), Contribution of Working Group I to the Sixth Assessment Report of the Intergovernmental Panel on Climate Change. Summary for Policymakers. Cambridge University Press, Cambridge, United Kingdom and New York, NY, USA, pp. 3–32, https://doi-org/10.1017/9781009157896.001.

Jackson JE (2015) Alternative Material Nest Boxes and Impacts on Nestling Physiology and Adult Behavior in the Eastern Bluebird (Sialia sialis).

Jankowski MD, Russell RE, Franson JC, Dusek RJ, Hines MK, Gregg M, Hofmeister EK (2014) Corticosterone Metabolite Concentrations in Greater Sage-Grouse Are Positively Associated With the Presence of Cattle Grazing. Rangel Ecol Manag 67: 237–246.

Johnson JL, Rollins D, Reyna KS (2012) What’s A Quail Worth? A Longitudinal Assessment of Quail Hunter Demographics, Attitudes, and Spending Habits in Texas. National Quail Symposium Proceedings 7: 294–299.

Jones SM, Lockhart TJ, Rose RW (2005) Adaptation of wild-caught Tasmanian devils (Sarcophilus harrisii) to captivity: evidence from physical parameters and plasma cortisol concentrations. Aust J of Zool 53: 339–344.

Khan MZ, Altmann J, Isani SS, Yu J (2002) A matter of time: evaluating the storage of fecal samples for steroid analysis. Gen Comp Endocrinol 128: 57–64.

Lattin CR, Breuner CW, Romero LM (2016) Does corticosterone regulate the onset of breeding in free-living birds?: The CORT-Flexibility Hypothesis and six potential mechanisms for priming corticosteroid function. Horm Behav 78: 107–120.

Leopold AS (1977) The California Quail. University of California Press, Berkeley, CA.

Lexen E, El-Bahr SM, Sommerfeld-Stur I, Palme R, Mostl E (2008) Monitoring the adrenocortical response to disturbances in sheep by measuring glucocorticoid metabolites in the faeces. Wien Tierarztl Monatsschr 95: 64–71.

Littin KE, Cockrem JF (2001) Individual variation in corticosterone secretion in laying hens. Br Poult Sci 42: 536–546.

Lynn SE, Stamplis TB, Barrington WT, Weida N, Hudak CA (2010) Food, stress, and reproduction: short-term fasting alters endocrine physiology and reproductive behavior in the zebra finch. Horm Behav 58: 214–222.

Martin JA, Applegate RD, Dailey TV, Downey M, Emmerich B, Hernández F, McConnell MM, Reyna KS, Rollins D, Ruzicka RE (2017) Translocation as a population restoration technique for northern bobwhites: a Review and Synthesis. Proceedings of the National Quail Symposium 8: 11–16.

McEwen BS, Wingfield JC (2003) The concept of allostasis in biology and biomedicine. Horm Behav 43: 2–15.

Messmann S, Bagu E, Robia C, Palme R (1999) Measurement of glucocorticoid metabolite concentrations in faeces of domestic livestock. J Vet Med A Physiol Pathol Clin Med 46: 621–631.

Millspaugh JJ, Washburn BE (2004) Use of fecal glucocorticoid metabolite measures in conservation biology research: considerations for application and interpretation. Gen Comp Endocrinol 138: 189–199.

Mohlman JL, Navara KJ, Sheriff MJ, Terhune TM, Martin JA (2020) Validation of a noninvasive technique to quantify stress in northern bobwhite (*Colinus virginianus*). Conserv Physiol 8: coaa026.

Möstl E, Rettenbacher S, Palme R (2005) Measurement of corticosterone metabolites in birds’ droppings: an analytical approach. Ann N Y Acad Sci 1046: 17–34.

National Research Council (2010) Guide for the Care and Use of Laboratory Animals, Eighth Edition. The National Academies Press, Washington, DC, USA.

Palme R (2019) Non-invasive measurement of glucocorticoids: advances and problems. Physiol Behav 199: 229–243.

Palme R, Fischer P, Schildorfer H, Ismail MN (1996) Excretion of infused 14C-steroid hormones via faeces and urine in domestic livestock. Anim Reprod Sci 43: 43–63.

Perez RM, Wilson DE, Gruen KD (2002) Survival and flight characteristics of captive-reared and wild Northern Bobwhites in South Texas. Proceedings of the National Quail Symposium 5: 81–85.

Pope M, Heekin P (2017) How Ecological Disturbances May Influence Mountain Quail in the Pacific Northwest. Proceedings of the National Quail Symposium 4.

Rabon JC, Nuñez CMV, Coates PS, Ricca MA, Johnson TN (2021) Ecological correlates of fecal corticosterone metabolites in female Greater Sage-Grouse (*Centrocercus urophasianus*). Can J Zool 99: 812–822.

Reyna KS, Newman WL (2018) Comparative analysis of behavioural response of captive-reared and wild-trapped Northern Bobwhites to simulated predator attacks. Avian Biol Res 11: 16–23.

Reyna KS, Whitt JG, Currier SA, SM SP, Rushing GT, Conley JT, Vandenberg CA, Moser EL (2020) California quail translocation from Idaho to Texas. The Quail Research Laboratory, College of Agricultural Sciences and Natural Resources, Texas A&M University Commerce.

Rich EL, Romero LM (2005) Exposure to chronic stress downregulates corticosterone responses to acute stressors. Am J Physiol Regul Integr Comp Physiol 288: R1628–R1636.

Romero LM, Remage-Healey L (2000) Daily and seasonal variation in response to stress in captive starlings (*Sturnus vulgaris*): corticosterone. Gen Comp Endocrinol 119: 52–59.

Romero ML, Butler LK (2007) Endocrinology of stress. Int J Comp Psychol 20: 89–95.

Rupp H (1999) Excess Catecholamine Syndrome: Pathophysiology and Therapy a. Ann N Y Acad Sci 881: 430–444.

Sapolsky RM, Romero LM, Munck AU (2000) How do glucocorticoids influence stress responses? Integrating permissive, suppressive, stimulatory and preparative actions. Endocr Rev 21: 55–89..

Scott JL, Hernández F, Brennan LA, Ballard BM, Janis M, Forrester ND (2013) Population demographics of translocated northern bobwhites on fragmented habitat. Wild Soc Bull 37: 168–176.

Şekercioğlu ÇH, Primack RB, Wormworth J (2012) The effects of climate change on tropical birds. Biol Conserv 148: 1–18.

Seneviratne S, Nicholls N, Easterling D, Goodess C, Kanae S, Kossin J, Luo Y, Marengo J, McInnes K, Rahimi M (2012) Changes in climate extremes and their impacts on the natural physical environment. In: Field CB, Barros VR, Stocker TF, Qin D, Dokken DJ, Ebi KL, Mastrandrea MD, Mach KJ, Plattner GK, Allen SK, et al., eds. Managing the Risks of Extreme Events and Disasters to Advance Climate Change Adaptation, A Special Report of Working Groups I and II of the Intergovernmental Panel on Climate Change (IPCC). Cambridge University Press, Cambridge, UK.

Sheriff MJ, Bosson CO, Krebs CJ, Boonstra R (2009) A non-invasive technique for analyzing fecal cortisol metabolites in snowshoe hares (*Lepus americanus*). J Comp Physiol B 179: 305–313.

Sheriff MJ, Dantzer B, Delehanty B, Palme R, Boonstra R (2011) Measuring stress in wildlife: techniques for quantifying glucocorticoids. Oecologia 166: 869–887.

Sheriff MJ, Krebs CJ, Boonstra R (2010) Assessing stress in animal populations: do fecal and plasma glucocorticoids tell the same story? Gen Comp Endocrinol 166: 614–619.

Shipley AA, Sheriff MJ, Pauli JN, Zuckerberg B (2019) Snow roosting reduces temperature-associated stress in a wintering bird. Oecologia 190: 309–321.

Shipley AA, Sheriff MJ, Pauli JN, Zuckerberg B (2022) Weather and land cover create a predictable “stress-scape” for a winter-adapted bird. Landsc Ecol 37: 779–793.

Sisson DC, II TMT, Palmer WE, Thackston R (2017) Contributions of translocation to northern bobwhite population recovery. Proceedings of the National Quail Symposium 8: 151– 159.

Sokół R, Koziatek-Sadłowska S (2020) Changes in the corticosterone level in tooting male black grouse (*Tetrao tetrix*) infected with *Eimeria* spp. Poultry science 99: 1306–1310.

Sokos CK, Birstas PK, Tsachalidis EP (2008) The aims of galliforms release and choice of techniques. Wildlife Biology 14: 412–422.

Sportsmen’s Alliance (2021) Economic Impact of Hunting and Shooting in 2020 Technical Report. Southwick Associates, Fernandina Beach, Florida, USA.

Stephens SE, Koons DN, Rotella JJ, Willey DW (2004) Effects of habitat fragmentation on avian nesting success: a review of the evidence at multiple spatial scales. Biol Conserv 115: 101–110.

Stephenson JA, Reese KP, Zager P, Heekin PE, Nelle PJ, Martens A (2011) Factors influencing survival of native and translocated mountain quail in Idaho and Washington. J Wildl Manage 75: 1315–1323.

Suedkamp Wells KM, Washburn BE, Millspaugh JJ, Ryan MR, Hubbard MW (2003) Effects of radio-transmitters on fecal glucocorticoid levels in captive Dickcissels. Condor 105: 805– 810.

Suorsa P, Huhta E, Nikula A, Nikinmaa M, Jäntti A, Helle H, Hakkarainen H (2003) Forest management is associated with physiological stress in an old–growth forest passerine. Proc R Soc Lond B Biol Sci 270: 963–969.

Tanner EP, Elmore RD, Fuhlendorf SD, Davis CA, Dahlgren DK, Orange JP (2017) Extreme climatic events constrain space use and survival of a ground-nesting bird. Glob Chang Biol 23: 1832–1846.

Tempel DJ, Gutiérrez RJ (2004) Factors related to fecal corticosterone levels in California spotted owls: implications for assessing chronic stress. Conserv Biol 18: 538–547.

Thiel D, Jenni-Eiermann S, Braunisch V, Palme R, Jenni L (2008) Ski tourism affects habitat use and evokes a physiological stress response in capercaillie *Tetrao urogallus*: a new methodological approach. J Appl Ecol 45: 845–853.

Thiel D, Jenni-Eiermann S, Palme R, Jenni L (2011) Winter tourism increases stress hormone levels in the Capercaillie *Tetrao urogallus*. Ibis 153: 122–133.

Touma C, Palme R (2005) Measuring fecal glucocorticoid metabolites in mammals and birds: the importance of validation. Ann N Y Acad Sci 1046: 54–74.

Touma C, Palme R, Sachser N (2004) Analyzing corticosterone metabolites in fecal samples of mice: a noninvasive technique to monitor stress hormones. Horm Behav 45: 10–22.

van Vliet HEJ, Stutchbury BJM, Newman AEM, Norris DR (2020) The impacts of agriculture on an obligate grassland bird of North America. Agric Ecosyst Environ 287: 106696.

Warwick H, Morris P, Walker D (2006) Survival and weight changes of hedgehogs (Erinaceus europaeus) translocated from the Hebrides to Mainland Scotland. Lutra 49: 89–102.

Washburn BE, Millspaugh JJ, Schulz JH, Jones SB, Mong T (2003) Using fecal glucocorticoids for stress assessment in mourning doves. Condor 105: 696–706.

Wasser SK, Hunt KE, Brown JL, Cooper K, Crockett CM, Bechert U, Millspaugh JJ, Larson S, Monfort SL (2000) A generalized fecal glucocorticoid assay for use in a diverse array of nondomestic mammalian and avian species. Gen Comp Endocrinol 120: 260–275.

Wasser SK, Risler L, Steiner RA (1988) Excreted steroids in primate feces over the menstrual cycle and pregnancy. Biol of Reprod 39: 862–872.

Whitt JG, Johnson JA, Reyna KS (2017) Two centuries of human-mediated gene flow in northern bobwhites. Wild Soc Bull 41: 639–648.

Wilsey C, Bateman B, Taylor L, Wu JX, LeBaron G, Shepherd R, Koseff C, Friedman S, Stone R (2019) Survival by degrees: 389 bird species on the brink. National Audubon Society: New York, NY, USA.

Wingfield JC, Pérez JH, Krause JS, Word KR, González-Gómez PL, Lisovski S, Chmura HE (2017) How birds cope physiologically and behaviourally with extreme climatic events. Philos Trans R Soc Lond B Biol Sci 372: 20160140.

Wingfield JC, Ramenofsky M (2011) Hormone-behavior interrelationships of birds in response to weather. Adv Study Behav 43: 93–188.

Wingfield JC, Romero LM (2011) Adrenocortical Responses to Stress and Their Modulation in Free-Living Vertebrates. In: Comprehensive Physiology. John Wiley & Sons, Ltd, pp 211–234.

Wingfield JC, Sapolsky RM (2003) Reproduction and resistance to stress: when and how. J Neuroendocrinol 15: 711–724.

Wszola LS, Gruber LF, Stuber EF, Messinger LN, Chizinski CJ, Fontaine JJ (2020) Use and expenditures on public access hunting lands. J Outdoor Recreat Tour 29: 100256.

Zar JH (1996) Biostatistical Analysis. Pearson. Upper Saddle River, NJ, USA.

